# Virulence evolution: thinking outside of the host

**DOI:** 10.1101/2024.05.23.595559

**Authors:** Luís M. Silva, Jacob C. Koella

**Affiliations:** Institute of Biology, University of Neuchâtel, Neuchâtel, Switzerland; Department of Zoology, The University of British Columbia, Vancouver, Canada

**Keywords:** pathogen evolution, virulence, transmission, infectivity, infection severity, microsporidia

## Abstract

The main theory of the evolution of virulence relies on a trade-off between virulence and transmission rate. However, it has been difficult to measure the required trade-off. A recent transmission decomposition framework explains that this might be partly due to a lack of information about the parasite’s survival in the environment outside its hosts, where the parasite finds itself during transmission to its next host. In this study, we used parasite lines of the microsporidian *Vavraia culicis* with varying levels of virulence upon infecting their host, the mosquito *Anopheles gambiae*, to explore the interaction between parasite-driven virulence within its host and its survival outside of the host. The parasite lines with greater virulence and growth within their hosts had a cost in their intrinsic ability to withstand the environment, irrespective of temperature. These results underscore the importance of considering the full context of transmission and other parasite fitness traits in studying and predicting the evolution and spread of infectious diseases.

## Introduction

Parasite fitness depends on successful transmission, which often requires a delicate balance between replicating within the host and enduring the environment among hosts (Silva *et al*. 2025a). Classical evolutionary theory suggests parasite replication, while necessary for producing transmissible stages, is constrained by the harm it causes the host, i.e., virulence (Anderson & May 1982; Jensen *et al*. 2006; De Roode *et al*. 2008). An increase in virulence caused by too much host damage may reduce host survival or contact rates, ultimately limiting the opportunities for parasite transmission. These ideas were compiled by Anderson and May (1982) and formed the foundation for most epidemiological, clinical, and conservation disease control strategies (Alizon *et al*. 2009; Anderson & May 1982). However, the evolutionary constraint imposed by this theory may be relaxed for parasites that persist in the environment after being shed.

This decoupling of transmission from host survival is central to two prominent theories in virulence evolution: the obligate killer and the Curse of the Pharaoh hypotheses. The former states that when transmission occurs only upon or after host death, as is the case for some bacteriophages or spore-forming parasites, selection may favour high virulence to maximize replication and ensure the timely release of the infectious stages (Ben-Ami 2017; Ebert & Weisser 1997; Redman *et al*. 2016). In such cases, killing the host is not a cost but a necessary step for transmission. The Curse of the Pharaoh hypothesis predicts that long-lived infective stages in the external environment reduce the cost of killing the host, thereby selecting for higher virulence (Bonhoeffer *et al*. 1996; Gandon 1998; Rafaluk-Mohr 2019). When transmission can still occur after host death, particularly when environmental conditions support the persistence of infectious stages, parasites are under less pressure to maintain host longevity. These theories underscore the importance of the external environment in shaping parasite evolution. Indeed, most parasites must endure a phase outside their host, where they are exposed to diverse abiotic and biotic factors (Silva *et al*. 2025a; Turner *et al*. 2021). Therefore, the nature and duration of this among-hosts phase can drive the evolution of parasite life-history traits, potentially favouring more virulent strategies in specific ecological contexts.

However, a meta-analysis performed by Rafaluk-Mohr (2019) showed that the relationship between virulence and environmental persistence is not as clear-cut as initially thought (Rafaluk-Mohr 2019). While specific taxa, such as viruses, showed a trade-off between the two, others, as fungi or bacteria, showed a positive association.

Hence, in this study, we sought to better understand the relationship between virulence, parasite transmission, and environmental survival. For this, we used the microsporidian *Vavraia culicis*, a generalist parasite of several dipteran species (Zeferino & Koella 2024) that contains an environmental stage among hosts (Silva *et al*. 2025c). We had earlier selected lines of *V. culicis* to transmit only upon the death of their host, the mosquito *Anopheles gambiae* (Silva & Koella 2025), resembling the obligate killer parasite strategy, and had established two selection regimes: i) *Early*, where the parasites within the first third of the mosquitoes to die were selected and used to infect the next generation of naive mosquitoes (i.e., parasites that killed the host under 7 days); or ii) *Late*, where the parasites from the last third of the mosquitoes to die were used to infect the next generation of mosquitoes (i.e., parasites that killed the host after 20 days). It is noteworthy to state that the selection for early and late killing also selected for shorter and longer times within the host, respectively. After seven passages, early and late performance within the host was assessed, and compared to an unselected parasite population, which grew parallel to the selection experiment: *Stock. Late* parasites shortened their life cycle, increased virulence (measured as maximum hazard of the death curves; hosts infected with *Early, Late* and *Stock* lived on average 18, 20 and 21 days, respectively) and replicated more rapidly (Silva & Koella 2025) due to more efficient iron sequestration and usage (Silva *et al*. 2025b) than either selected *Early* or unselected *Stock*. Furthermore, in response to infection by Late parasites, the host shift its investment from immunity to earlier fecundity (Silva 2024; Silva & Koella 2025).

In this experiment, we allowed the *V. culicis* lines to replicate within a naive population of mosquitoes for 20 days. After spore production, we prepared ten aliquots from each parasite treatment and replicate. Each aliquot contained the standard dose of 10,000 spores per larva in one millilitre of distilled water, to which an antimycotic-antibiotic cocktail was added to prevent undesired microbial growth during the duration of the experiment. Aliquots were randomly stored at one of two temperatures, 4 °C or 20 °C. Then, immediately after (time point 0 days), 45 and 90 days later, the stored spores were used to infect naive populations of *A. gambiae* mosquitoes. The number of infected individuals (infectivity) and the load of the infected individuals (infection severity) were then assessed for each parasite treatment and replicate.

We found that the *V. culicis* lines that are better adapted to the host, with high virulence and cunning exploitation, are less able to survive outside it. Although *V. culicis* transmission heavily depends on its environmental stage (spores), our findings show an inverse relationship between parasite replication and virulence evolution, in contrast to previously presented theories. Altogether, our findings highlight the importance of the environment and host-parasite traits in understanding virulence evolution and parasite transmission.

## Materials and Methods

### Experimental model

We used the Kisumu strain of the mosquito host *Anopheles gambiae (s*.*s*.) and lines of the microsporidian parasite *Vavraia culicis floridensis* selected for seven generations: i) for early transmission and shorter time within the host or for late transmission and longer time within the host; ii) or lines taken from our unselected stock as a baseline reference (Silva & Koella 2025). Further details on the selection experiment and its maintenance are described in Silva and Koella (2025). Experiments and rearing were conducted at standard laboratory conditions (26 ± 1°C, 70 ± 5% relative humidity, and 12h light/dark).

### Preparation of parasite aliquots

Freshly hatched *A. gambiae* larvae were reared in groups of 50 per Petri dish (120 mm diameter x 17 mm) containing 80 ml of distilled water. They were fed daily with Tetramin Baby fish food according to their age: 0.04, 0.06, 0.08, 0.16, 0.32 and 0.6 mg/larva for 0, 1, 2, 3, 4 and 5 days or older, respectively. Two days after hatching, larvae were exposed to 10,000 spores of *V. culicis* per larva, from spores obtained parasites selected for early-transmission, late-transmission, or not selected (i.e., stock). Each treatment consisted of five replicate lines maintained in similar conditions. Upon pupation, individuals were transferred to cages (21 × 21 × 21cm) and left to emerge as adults. The adults were provided constant access to a 6% sucrose-water solution. Dead mosquitoes were collected daily after their death up to day 20 (when the experiment ended), when the remaining alive mosquitoes were also collected. The collected mosquitoes were put into 2ml microcentrifuge tubes with 1ml of distilled water, with at most 25 mosquitoes per tube, and kept at 4°C and in the dark. At the end of the collection, a stainless-steel bead (Ø 5 mm) was added to each of the tubes and the mosquitoes were homogenized with a Qiagen TissueLyser LT at a frequency of 30 Hz for two minutes. The resulting homogenate containing the parasite spores was transferred to a 15 ml falcon tube, and the spore concentration was estimated with a haemocytometer under a phase-contrast microscope (400x magnification). From each falcon tube, we prepared aliquots with a spore concentration of 500,000 spores/ml for the infections in the following experiment. The resulting aliquots contained spores in 1:1 distilled water and an antibiotic-antimycotic cocktail solution (Sigma-Aldrich A5955, containing 10,000 units penicillin, 10 mg streptomycin and 25 μg amphotericin per ml, filtered), which limits the growth of contaminating microbes without affecting microsporidia (Van Gool *et al*. 1994; Jaronski 1984; Visvesvara 2002; Visvesvara *et al*. 1999). A preliminary experiment was run to ensure that the antibiotic-antimycotic solution did not interfere with the production or development of the parasite’s spores. Aliquots were kept in the dark at 4 ± 0.5 °C or 20 ± 0.5 °C dark until they were used.

### Infectivity and infection severity experiment

To assay how time in the external environment and temperature of the environment affect infectivity, we infected mosquitoes (reared as described above) with one of the aliquots prepared 0, 45, or 90 days earlier (Fig. 1). We used each aliquot only once to avoid contamination. After pupation, mosquitoes were moved to individual 50 ml falcon tubes containing approximately 10 ml of distilled water (Dao *et al*. 2010), and at adult emergence, they were transferred to individual cups (5 cm Ø x 10 cm) with access to a 6% sugar solution and a humid filter paper. Ten days after emergence, the mosquitoes were frozen at −20°C for later spore counts. This time point was chosen as all mosquitoes infected with any of the parasite lines have started to produce spores (Silva & Koella 2025).

**Figure 1.**
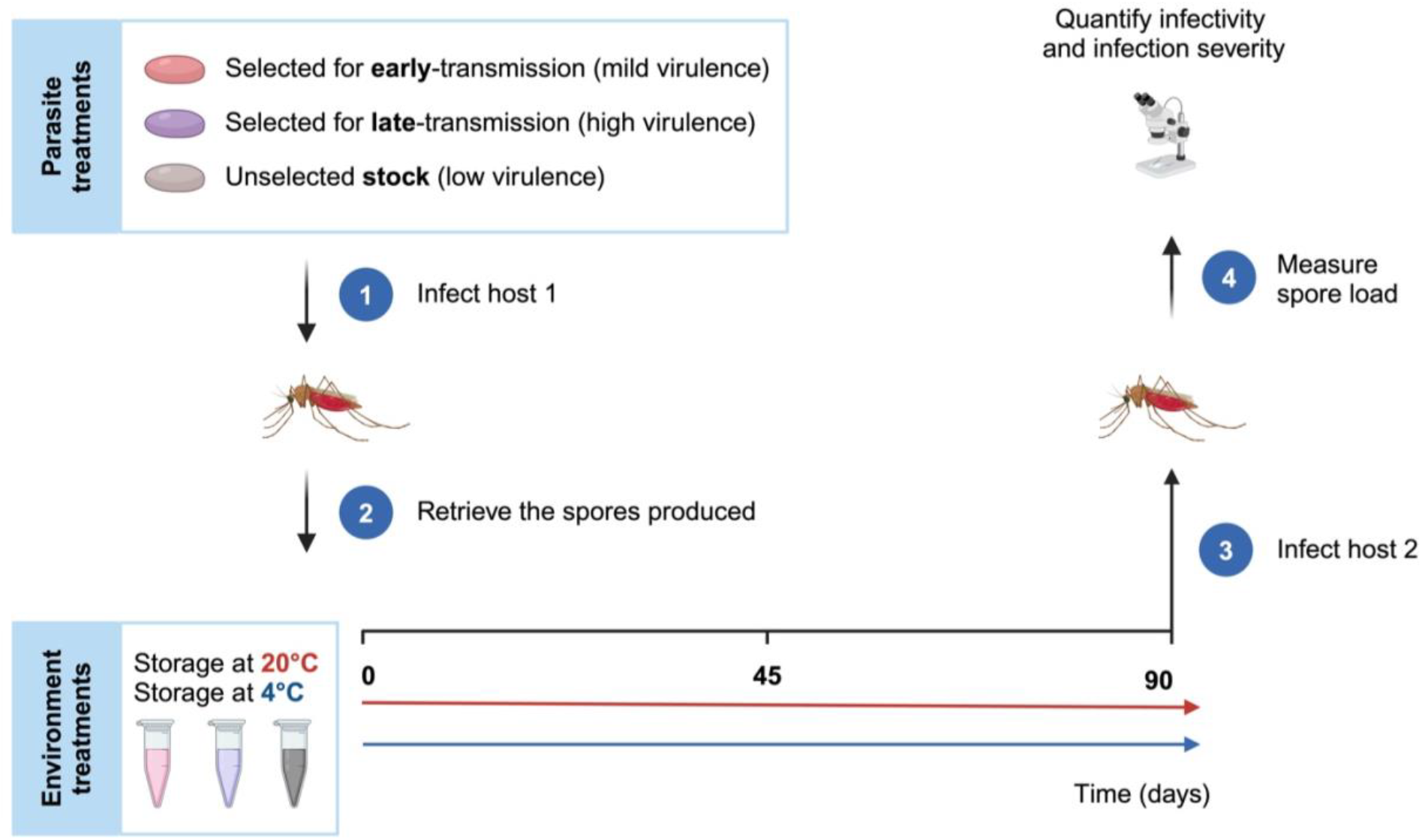
Experimental design. Summary illustration of the experimental design and respective workflow. In brief, different microsporidian parasite *V. culicis* lines were used to infect *A. gambiae* mosquitoes. The spores were stored at one of two temperatures (i.e., 4 °C or 20 °C) for 0-, 45- and 90-days post-spore-production and then used to infect naive *A. gambiae* mosquitoes. Spore load was then measured at day 10 of adulthood to assess the infectivity and infection severity of the different parasite lines. Figure generated with Biorender.com.

### Statistical analysis

The analyses were performed with R version 4.3.1 in RStudio version 3034.06.2+561. The packages “DHARMa” (Hartig & Hartig 2017), “car” (Fox *et al*. 2012), “lme4” (Bates *et al*. 2009) and “emmeans” (Lenth *et al*. 2019) were used to analyse the data. We used the packages “ggplot2” (Wickham 2016) and “scales” (Wickham *et al*. 2022) to plot the data and Biorender.com to assemble the figure panels.

Spore infectivity was measured as the proportion of infected individuals (Fig. 2a). Hence, we assessed differences in infectivity using a generalised linear mixed model with a binomial error, parasite selection, time in the external environment, temperature, and all interactions as explanatory variables.

**Figure 2.**
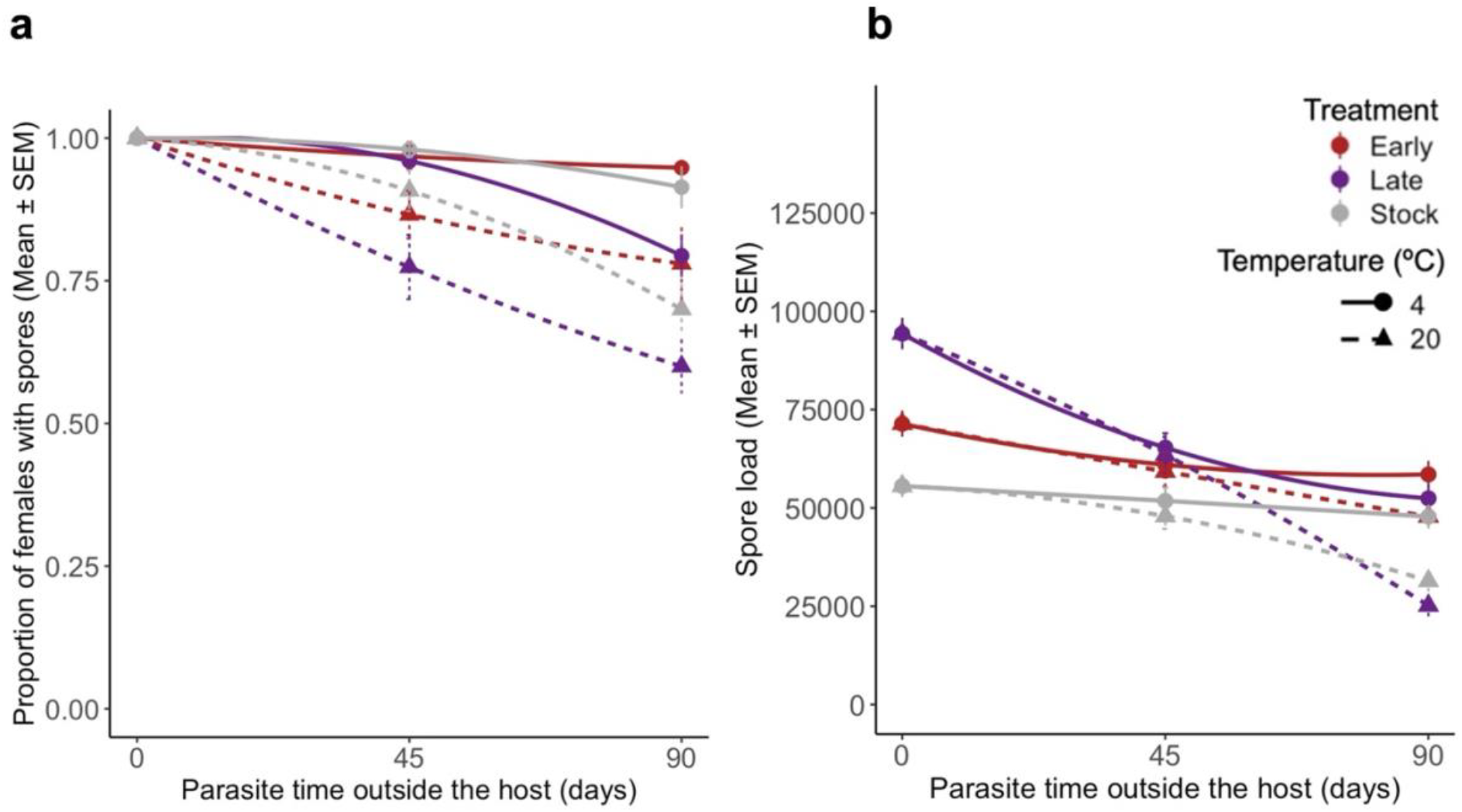
Parasite performance in a new host. **(a)** Infectivity of the spores by regime, time, and storage temperature. The latter was defined as the proportion of successful infections by day 10 of adulthood. Each mean consists of 150 females per treatment, time and temperature, split through five replicates. **(b)** Infection severity of the females carrying spores for the same time-point, as the number of spores per female. The sample size is the following for 4°C and 20°C, respectively. Day 0: All treatments consisted of 150 females. Day 45: Early with 145 and 130, Late with 144 and 116, Stock with 147 and 136. Day 90: Early 142 and 117, Late 119 and 90, Stock with 137 and 105. For further details on the statistics, see Table S1.

The selection line was included as a random factor. Infection severity was defined as the spore load of individuals containing spores (Fig. 2b). Infection severity was analysed similarly to the infectivity analyses, using a linear mixed model with a Gaussian error, the same explanatory variables, and replicate as a random factor.

Following the framework developed by Silva *et al*. (2025), we then quantified differences in spore quality (*Q*_*p*_) and environmental effect (*Q*_*e*_) for spore infectivity and infection severity (Silva *et al*. 2025a) (Fig. 3ab). *Q*_*p*_ for infectivity was calculated as the difference within each selection line of infectivity (proportion of individuals with spores) at 0 days and 90 days (Fig. 3a). *Q*_*p*_ for infection severity was calculated as the difference of spore load obtained after 0 days and after 90 days (Fig. 3b). As our standard environment we considered 4°C, the temperature the spores have been stored to for several years before this experiment. In both cases, we used a linear mixed model with parasite treatment as the explanatory variable and the replicate as a random factor. *Q*_*e*_ for infectivity was calculated as the difference of infectivity between 4°C and 20°C for the time-point of 90 days, where we observe the most striking difference due to temperature (Fig. 3b). *Q*_*e*_ for infection severity was the difference of the average spore load obtained after 90 days between parasite kept at 4 °C and 20°C days (Fig. 3b). We analysed both using a linear mixed model with parasite treatment as the explanatory variable and the replicate as a random factor.

**Figure 3.**
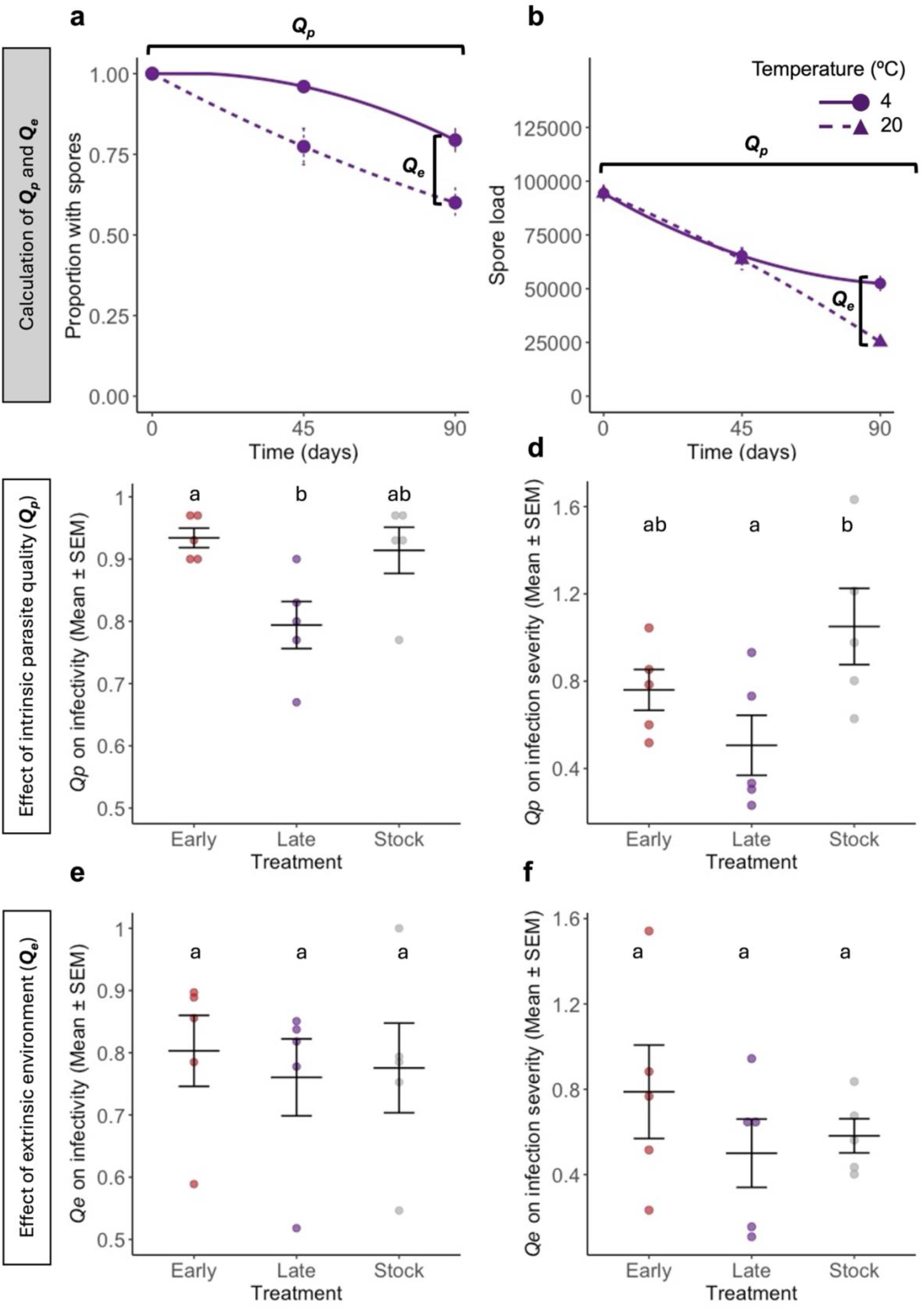
Calculation of the *Qp* and *Qe* for infectivity and infection severity. Parasite quality (*Q*_*p*_) and environment (*Q*_*e*_) effects on infectivity **(a)** and infection severity **(b)** were estimated, as demonstrated in the figures. In these figures, we demonstrate this rationale using the late-selected spores’ infectivity and infection severity curves per environmental temperature and by time post-spore production presented in Figure 1. *Q*_*p*_ was measured as the mean difference in the proportion of females with spores/spore load between infection with 90-days and 0-days old spores at 4° C. *Q*_*e*_, as the effect of environment temperature in this model, was calculated by subtracting the mean proportion of females with spores/spore load after exposure to spores kept at 20°C for 90 days from the mean proportion of females with spores/spore load at 4°C for 90 days. The resulting means of each replicate from each treatment were then used in Figures 2c-e. The effects of parasite selection on the parasite spore quality (*Q*_*p*_) and environmental temperature (*Q*_*e*_) for **(ce)** infectivity and **(df)** infection severity (i.e., spore load). Each data point represents the mean value of one of the five replicated selection lines. For further details on the statistical analyses, please see Table S2. *Q*_*p*_ was significantly linked to the average virulence of a selection line (Figs. 4ab; Table S3), with more virulent lines being less able to infect a new host (χ2 = 13.00; *df* = 2; *p <* 0.001; Table S3) and producing fewer spores in the new host (χ2 = 4.58; *df* = 2; *p =* 0.032; Table S3) after a long time in the external environment. However, the environmental sensitivity component, *Q*_*e*_, was not linked to virulence (Fig. 4cd; Table S3).

We used the maximum hazards from the survival curves generated and calculated by Silva and Koella (2025) as proxies for infection virulence in the first host. Maximum hazard values work as a proxy for the maximum death rate for each parasite treatment. We related virulence to infectivity by assessing whether the virulence of each selection line explained the differences in *Q*_*p*_ or *Q*_*e*_ for infectivity or infection severity. This was done using linear models, where the dependent and the independent variables were transformed by adding one and taking the logarithm. The curves in Fig. 4a-d were calculated using the respective lm function.

**Figure 4.**
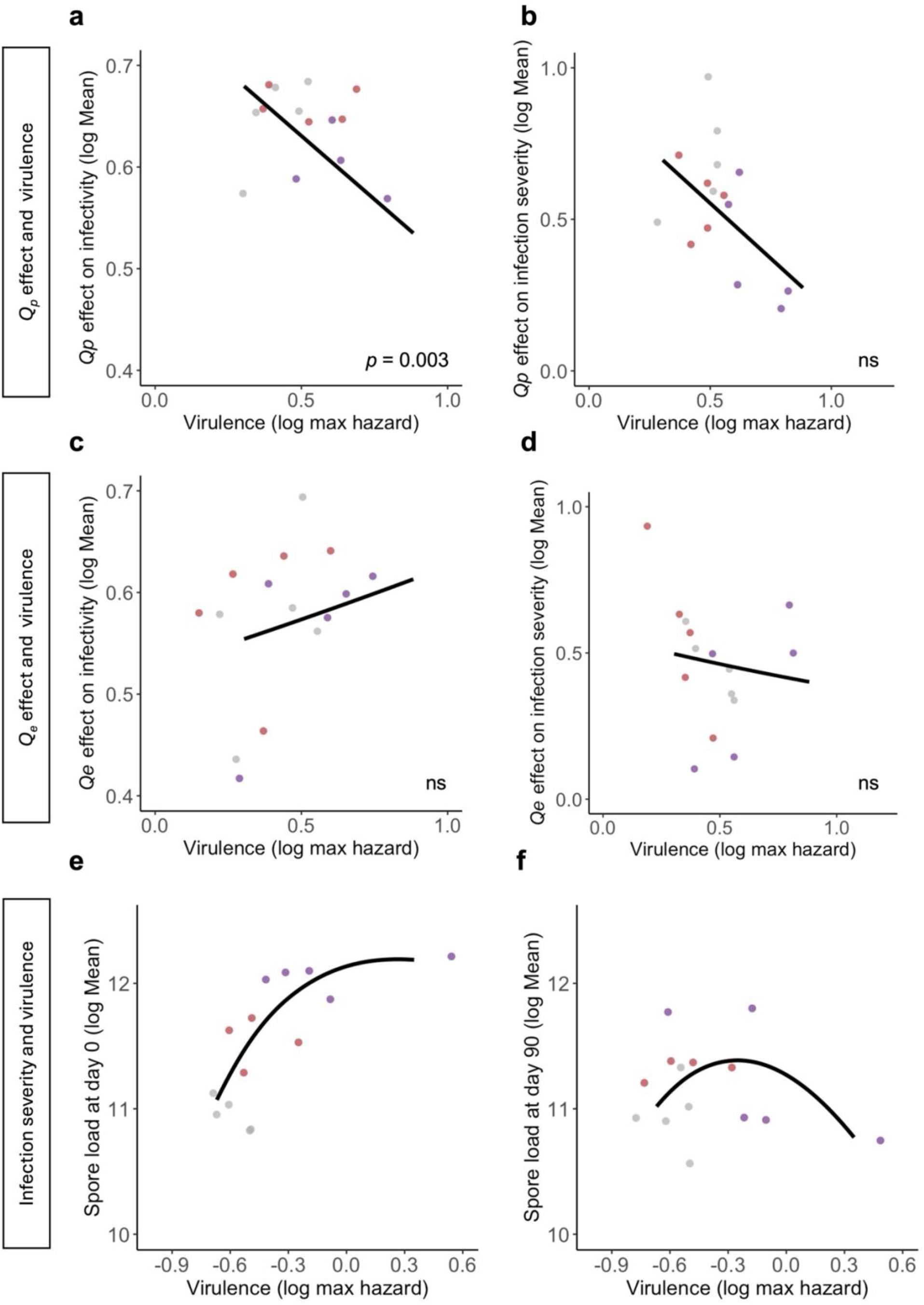
Virulence-transmission decomposition. Parasite’s virulence in the first host in relationship to *Q*_*p*_ *and Q*_*e*_ effect on infectivity **(ac)** and infection severity **(bd)** in a new host, respectively. Each point represents the average values of a replicated selection line. The curves show the result of the linear model. Parasite burden in a new host is caused after a time between hosts of 0 **(e)** and 90 days postproduction **(f)**. In these last two plots, a smooth line with the “loess” method was generated for ease of visualisation. See Table S3 for more details on the statistical analyses.

## Results

The direction of selection affected the parasite’s ability to establish an infection in a new host, with stock and early-selected parasites being more infective than late-selected parasites (χ2 = 33.00, *df* = 2, *p* < 0.001; Fig. 2a; Table S1). The longer the spores were kept in the external environment, the less infective they were (χ2 = 77.56, *df* = 1, *p* < 0.001; Table S1). If they were kept in the external environment at 20°C, they were less infective than if they were kept at 4°C (χ2 = 166.85, *df* = 1, *p* < 0.001; Fig. 2a; Table S1). None of the interactions between selection treatment, temperature and duration in the external environment was significant. A similar pattern was observed for spore load (infection severity) of established infections (Fig. 2b; Table S1) (selection: χ2 = 135.31, *df* = 2, *p* < 0.001; duration in the external environment: χ2 = 14.23, *df* = 2, *p* < 0.001; temperature: χ2 = 4.46; *df* = 1; *p =* 0.035; Table S1). The effect of the duration in the external environment (interaction selection * duration: χ2 = 45.51, *df* = 2, *p* < 0.001; Table S1) and of temperature (interaction selection * temperature: χ2 = 7.35; *df* = 1; *p =* 0.025; Table S1) was more pronounced for parasites selected for late transmission than for early-selected or unselected parasites.

While the component of intrinsic quality, *Q*_*p*_, differed across parasite treatments for both infectivity and infection severity (infectivity: χ2 = 11.286, *df* = 2, *p* = 0.004; infection severity: χ2 = 9.022, *df* = 2, *p* = 0.011 Figure 3cd; Table S2), the component of environmental sensitivity, or *Q*_*e*_, did not (Figure 3ef; Table S4). This finding suggests the parasites differ in their quality and performance, irrespective of the environmental pressure (i.e., temperature).

## Discussion

Most parasites spend some of their life cycle outside the host when transmitting from one host to the next. While this phase of parasitic life cycles is often ignored in evolutionary ideas, it has been considerably studied and described concerning viruses (Kormuth *et al*. 2019; Kotwal & Cannon 2014) and parasites with complex life cycles (Bradley *et al*. 2018; Davies *et al*. 2001). Here, we suggest that integrating and relating within-to outside-host parasite evolution may lead to important insights, mainly if the efficacy of transmission during the period spent outside the host trades off with growth and virulence within the host. A standard prediction, for example, is that selecting for later transmission would result in slower parasite growth within the host and lower virulence. However, an experiment with the artificial selection lines of the parasite *V. culicis* selected for early or late transmission showed the opposite pattern (Silva & Koella 2025). As discussed by the authors, this unexpected result could arise from the fact that the experimental design eliminated any costs of virulence during the transmission phase in the external environment between hosts. This experimental design favoured the selection of parasites that invest in exploitation and survival within the host rather than outside of it (Silva & Koella 2025). This study’s findings corroborate this idea and highlight the importance of considering transmission as a whole in parasite evolution (Boldin & Kisdi 2012; Silva *et al*. 2025a; Visher *et al*. 2021).

First, the longer the spores were kept outside of a host, the less able they were to infect a new host (Fig. 2a). This result was particularly notable in the lines with the highest growth within the host (that is, the lines selected for late transmission), indicating a trade-off between growth and virulence within the host and survival outside of the host. Other studies have suggested a trade-off between virulence and survival either outside of the host or between hosts (Davies *et al*. 2001; Dearsly *et al*. 1990; Leggett *et al*. 2014), and indeed, adaptation to one environment (e.g., within the host) is generally expected to incur costs in other environments (e.g. outside of hosts) (Caraco & Wang 2008). Similar results have been observed in viral studies when viruses are subjected to different environments throughout their transmission cycle (Bhardwaj & Agrawal 2020; Brandon Ogbunugafor *et al*. 2013; Wasik *et al*. 2023). Infection severity in the next host (that is, spore load) showed similar results from infectivity (Fig. 2b). This result suggests that the mortality of spores leads to less effective infectivity, which in turn results in a smaller dose of infection and thus (as observed in other parasites) slower within-host growth later on (Acuña-Hidalgo *et al*. 2022; Duneau *et al*. 2017; Torres *et al*. 2016). Additionally, since infection severity decreases more rapidly than infectivity between 45- and 90-day-old spores in the external environment, time outside the host affects not only the survival of the infective spores but also their infection potential. This suggests that an additional process is at play. One possibility is that the resources within a spore may be consumed or degraded over time, which could delay its growth upon entering the host. Such a process, called growth arrest, has not been described for microsporidia but is common in bacteria (Bergkessel *et al*. 2016; Hobbie & Hobbie 2013). It helps the bacteria meet their energetic and metabolic requirements before restarting replication when cells become more susceptible to environmental stressors (Moreno-Gámez *et al*. 2020). Another hypothesis is that parasites with free-living stages invest in processes and metabolic changes that allow them to subsist harmful abiotic factors (e.g., ultraviolet radiation and chemical exposure) for a considerable time. Such changes in metabolism require time and the suitable precursors and signalling to be reversed, which might hinder their instantaneous growth within a new host (Caraco & Wang 2008; Leblanc & Lefebvre 1984).

Second, the decrease of infection success in a new host with time spent outside of a host was stronger when the spores were kept at 20°C than when they were kept at 4°C (Fig. 1). Reduced spore survival at higher temperatures has been described for other aquatic microsporidia species, which thrive at lower temperatures (Becnel & Weiss 2014). In addition to their natural ecology, the parasite used in our experiment has been stored at cold temperatures, for in our lab, it is stored in the fridge (i.e., at 4 °C). Again, we cannot discard the fact that adaptation to a colder environment might have happened. Nevertheless, literature on aquatic microsporidia suggests they naturally survive better at lower temperatures (Becnel & Weiss 2014).

Third, by decomposing the parasite’s transmission success across different selection lines, an unselected parasite population, and temperatures, we were able to disentangle the extrinsic effect of the environment (*Q*_*e*_) and the parasite’s intrinsic quality (*Q*_*p*_) on infectivity and infection severity (Fig. 3) (Silva *et al*. 2025a). This allowed us to describe how each of these components relates to parasite’s virulence within the host. While *Q*_*e*_ effect on infectivity or infection severity seems unrelated to any treatment, *Q*_*p*_ was. Thus, late-selected spores were less resistant to time outside the host than early-selected and control ones. Still, this resilience was intrinsic to the parasite and not affected by temperature (Fig. 3). Furthermore, *Q*_*p*_, but not *Q*_*e*_ for both measures of transmission were negatively correlated with virulence (Fig. 4ab), suggesting a trade-off between virulence and parasite endurance in the external environment and corroborating that some microbes have a trade-off between reproduction and survival of their offspring generations. Such a hypothesis has been highly debated in the literature (Bonhoeffer *et al*. 1996; Caraco & Wang 2008; Rafaluk-Mohr 2019; Silva *et al*. 2025a), as there are examples of parasites with high virulence and low persistence in the outside environment, as well as very high (Rafaluk-Mohr 2019). The last strategy is named the Curse of the Pharaoh hypothesis, which states that virulence and persistence in the environment might be disentangled (Bonhoeffer *et al*. 1996) in specific ecological conditions. The parasite’s prevalence in an environment could be secured in this scenario, even in fluctuating host numbers. Hence, depending on the ecological circumstances of the host-parasite relationship, we can observe both patterns in nature (Rafaluk-Mohr 2019). Nevertheless, in our case, and likely most parasites, *V. culicis* is a self-sustaining parasite that can colonise multiple hosts and prevails in different host populations (Becnel & Weiss 2014; Zeferino & Koella 2024), which suggests a considerable genetic diversity that would not justify investing in the strategy described by the Curse of the Pharaoh (Bonhoeffer *et al*. 1996). In this case, evolution should favour pleiotropy constraints between virulence and persistence outside of the host, such as the negative relationship observed in this study (Fig. 4). It is important to note that the experiment was conducted using a single host genotype, and as such, the observed effects of parasite intrinsic quality (*Qp*) on infectivity and infection severity are contingent on this genetic background. Future work incorporating host genetic variation—and ultimately host–parasite genotype interactions—will be essential to assess the generality and ecological relevance of these findings.

Overall, our findings emphasize the importance of considering both ecology and the whole parasitic transmission cycle when studying parasite and virulence evolution (Carlsson & Råberg 2024; Silva *et al*. 2025a). By quantifying parasite survival to different environmental parameters, we could better explain the evolution of virulence in this system. Moreover, by using the transmission framework, we could associate this cost in parasite longevity outside of the host to the intrinsic properties of the parasite. The assumptions and conclusions of this study are of general importance. They can be applied to several taxa, as they reflect most of the parasite’s transmission cycles and consequent evolutionary paths. In fact, when we glance over the relationship between infection severity in a new host and virulence, we observe one of the advantages of having the full picture over transmission. While infection severity with freshly produced spores has a linear relationship with virulence (Fig. 4e), the same trait with 90-day-old spores produces a concave shape (Fig. 4f), as expected from the virulence-transmission trade-off theory. Hence, we believe the approach taken in this study undeniably reflects the complex network between transmission and other parasite fitness traits, as well as the limitations and possible negative consequences of only looking at within-host parasite fitness. The study and dissection of this network and their inherent trade-offs have major ramifications for studying disease ecology and parasite evolution. We can only better predict disease spread and plan consequent control strategies by understanding the latter. For instance, the findings in this study alert us that certain social behaviours (e.g., crowding) or social settings directly affecting susceptible host density might reduce the parasite time outside of the host, which in turn might inadvertently select for more virulent strains within a new host.

## Supplementary information

**Table S1.** Parasite infectivity and infection severity in a new host.

| **Tested effect** |  |  |  |
| --- | --- | --- | --- |
| **Model 1a - Proportion of females with spores** | **$\chi^2$** | ***df*** | ***p*** |
| Treatment | 32.996 | 2 | **< 0.001** |
| Temperature | 77.557 | 1 | **< 0.001** |
| Time | 166.849 | 1 | **< 0.001** |
| Treatment: Temperature | 1.011 | 2 | 0.603 |
| Treatment: Time | 3.446 | 2 | 0.179 |
| Temperature: Time | 0.433 | 1 | 0.511 |
| Treatment: Temperature: Time | 1.867 | 2 | 0.393 |
| **Model 1b - Spore load** | **$\chi^2$** | ***df*** | ***p*** |
| Treatment | 135.307 | 2 | **< 0.001** |
| Temperature | 0.002 | 1 | 0.961 |
| Time | 14.228 | 1 | **< 0.001** |
| Treatment: Temperature | 0.814 | 2 | 0.666 |
| Treatment: Time | 45.506 | 2 | **< 0.001** |
| Temperature: Time | 4.459 | 1 | **0.035** |
| Treatment: Temperature: Time | 7.352 | 2 | **0.025** |

**Table S2.** *Qp* and *Qe* on infectivity and infection severity.

| **Tested effect** |  |  |  |
| --- | --- | --- | --- |
| **Model 2c - $Qp$ on infectivity** | **$\chi^2$** | **$df$** | **$p$** |
| Treatment | 11.286 | 2 | **0.004** |
| Multiple comparisons for Model 2a |  | **$t$-ratio** | **$p$** |
| Early - Late |  | 3.106 | **0.035** |
| Early - Stock |  | 0.444 | 0.900 |
| Late - Stock |  | -2.662 | 0.067 |
| **Model 2d - $Qp$ on infection severity** | **$\chi^2$** | **$df$** | **$p$** |
| Treatment | 9.022 | 2 | **0.011** |
| Multiple comparisons for Model 2b |  | **$t$-ratio** | **$p$** |
| Early - Late |  | 1.884 | 0.205 |
| Early - Stock |  | -1.083 | 0.550 |
| Late - Stock |  | -2.968 | **0.043** |
| **Model 2e - $Qe$ on infectivity** | **$\chi^2$** | **$df$** | **$p$** |
| Treatment | 0.247 | 2 | 0.884 |
| **Model 2f - $Qe$ on infection severity** | **$\chi^2$** | **$df$** | **$p$** |
| Treatment | 1.809 | 2 | 0.405 |

**Table S3.** Virulence-transmission decomposition.

| **Tested effect** |  |  |  |
| --- | --- | --- | --- |
| **Model 4a - $Qp$ on infectivity vs. virulence** | **F** | ***df*** | ***p*** |
| log(Maximum hazard +1) | 12.934 | 1 | < **0.001** |
| **Model 4b - $Qp$ on infection severity vs. virulence** | **F** | ***df*** | ***p*** |
| log(Maximum hazard +1) | 4.351 | 1 | 0.057 |
| **Model 4c - $Qe$ on infectivity vs. virulence** | **F** | ***df*** | ***p*** |
| log(Maximum hazard +1) | 0.464 | 1 | 0.508 |
| **Model 4d - $Qe$ on infection severity vs. virulence** | **F** | ***df*** | ***p*** |
| log(Maximum hazard +1) | 0.155 | 1 | 0.701 |

## Notes

### Competing Interest Statement

The authors have declared no competing interest.

### Summary of Updates

Introduction and few other sentences were changed after revision process.

### Funding

- Basler Stiftung fu&#776;r biologische Forschung
- Swiss National Science Foundation: 310030_192786

